# AI-enabled reconstruction of 3D spatial multi-omics at single-cell resolution

**DOI:** 10.64898/2026.07.09.737490

**Authors:** Zhikang Wang, Yunzhi Yan, Xinwang Yang, Daoliang Zhang, Chuangyi Han, Qi Zou, Yixuan Du, Zheqi Hu, Zhiyuan Yuan

## Abstract

Three-dimensional (3D) spatial multi-omics provides unparalleled insights into biological activities, yet remains technically prohibitive. Here, we introduce Histo3D-MO, a hybrid experimental-computational pipeline for reconstructing single-cell-resolution 3D spatial multi-omics maps. Notably, Histo3D-MO integrates sparse, omics-disjoint spatial measurements with dense Hematoxylin and Eosin (H&E) histology through SPatial multi-Omics from h&E imaGEs (SPONGE), achieving cell-level 3D mapping across multiple omics layers. Validated using held-out slices, SPONGE substantially outperforms existing omics prediction methods. We further developed an algorithmic suite for 3D cell-type propagation and tissue-domain annotation, enabling whole-volume characterization of the tumor microenvironment. Applied to the in-house hepatocellular carcinoma data, Histo3D-MO revealed spatially organized patterns of translation efficiency, volumetric decoupling between malignant cells and monocytes, and depth-associated monocyte differentiation trajectories. Together, these results establish Histo3D-MO as a scalable framework for reconstructing single-cell-resolution 3D spatial multi-omics and interrogating tissue organization across complex biological systems.

## 1. Main

Spatial single-omics sequencing has demonstrated advantages in resolving complex biological systems by capturing molecular information within their native spatial context. As biological questions increasingly shift toward understanding higher-order tissue organization and functional heterogeneity, the prevailing analytical paradigm is evolving toward expanded molecular coverage (multi-omics)^1^ and spatial dimension (three-dimensional, 3D)^2^.

Recent advancements in spatial multi-omics^1,3^ enable simultaneous profiling of diverse molecular layers from single tissue sections. They are confronted with trade-offs between molecular coverage, regions of interest, spatial resolution, and the complexity of the experimental workflow^4^. 3D spatial omics sequencing is a more challenging task. Existing techniques can be categorized as serial-sectioning workflows and non-disruptive volumetric imaging^5^. The former requires iterative physical sectioning followed by individual profiling, with resources and costs scaling prohibitively with specimen volume. Volumetric techniques, though non-destructive, are limited by shallow penetration depth and restricted sequencing coverage (e.g., Deep-STARmap^2^: ∼200 μm depth, ∼1,017 genes). Consequently, 3D spatial multi-omics sequencing, requiring both volumetric continuity and multi-layer molecular profiling, remains a largely unexplored grand challenge.

Emerging evidence indicates that tissue morphology is intrinsically coupled with molecular expression profiles^6^. Inspired by this, we propose **Histo3D-MO** (Histology-informed 3D Spatial Multi-Omics), a **hybrid pipeline** that integrates experimental-computational strategies to enable the first **single-cell-resolution 3D spatial mapping across multiple molecular layers (≥2)**. It essentially takes the morphology features as the anchor to transfer the distinct omics features from only a few slices to the whole tissue volume (framed by multiple slices). By leveraging sparse spatial single-omics data alongside dense Hematoxylin & Eosin (H&E) histology, Histo3D-MO overcomes the inherent scalability and technique bottlenecks of multi-omics and 3D omics sequencing, thereby advancing **economical** 3D spatial multi-omics mapping. As its core, the proposed **SPONGE** (SPatial multi-Omics from h&e imaGEs) leverages H&E-derived morphological features to enable flexible multi-omics **diagonal integration**. Additionally, coupled with our designed cell-type and spatial-domain inference strategies, this framework facilitates integrated 3D tissue analysis.

An overview of Histo3D-MO is illustrated in Fig. 1a. We use our in-house Hepatocellular Carcinoma (HCC) dataset to demonstrate and evaluate the proposed framework. The workflow commenced with a dense sectioning of a tissue into *N* (*N*=100) serial slices (designated ID1∼ID100). Then, a minimal subset of *n* slices (*n*<< *N, n* =3) (ID1, ID10, ID100) were selected as “anchor” slices, with each implemented with single-cell–resolution spatial multi-omics sequencing, i.e., large-panel spatial transcriptomics and spatial proteomics. Two non-overlapping transcriptomic panels (indicated as Omics 1 and Omics 2) were assigned to slices ID1 and ID10, respectively, whereas the proteomic modality (Omics 3) was assigned to slice ID100, simulating our designed paradigm (sparse spatial single-omics data) and ensuring subsequent evaluation of generated omics features (diagonal integration). The remaining (*N – n*) intermediate slices are profiled using H&E imaging alone, providing a high-resolution structural scaffold for subsequent computational integration.

**Fig. 1.**
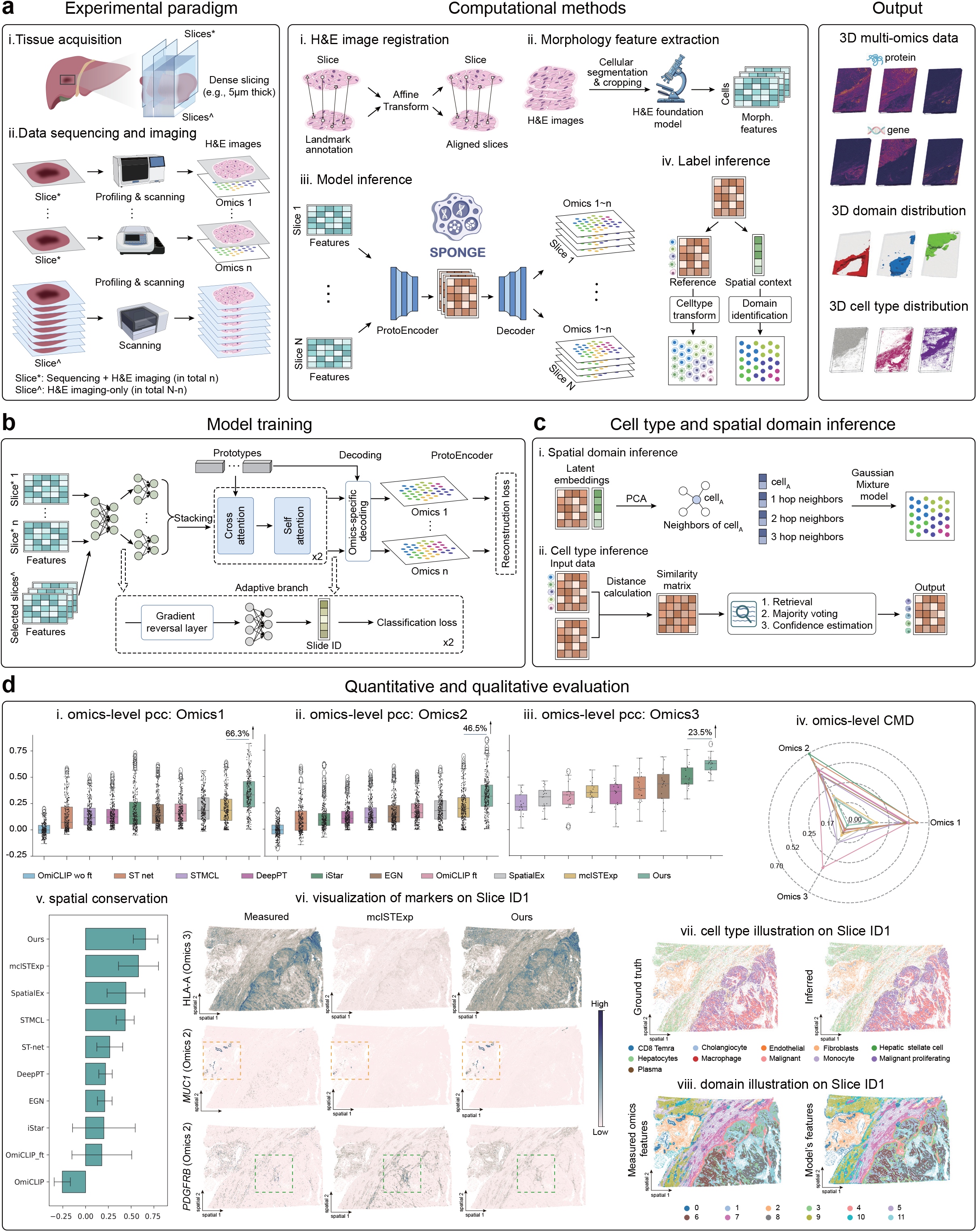
Overview of Histo3D-MO and performance of the SPONGE framework. a, Schematic illustration of the Histo3D-MO workflow, comprising the experimental paradigm (left), computational methods (middle), and model outputs (right). b, Training strategy of the SPONGE network, where both sequenced and imaging-only tissue slices are jointly incorporated to optimize model performance. c, Spatial domain (top) and cell type (bottom) inference strategies. d, Comprehensive evaluation of the trained SPONGE model, including both quantitative and qualitative assessments of omics prediction, spatial domain identification, and cell type inference. Panels **i–v** summarize quantitative comparisons across multiple omics prediction methods using diverse evaluation metrics. **vi**, Representative predictions from the second-best performing method for two marker genes and one protein. **vii**, Cell type inference results based on measured omics features (left) and cross-slice transfer from slices ID10 and ID100 to ID1 (right). **viii**, Spatial domain identification using measured omics data (left) compared with results from the proposed strategy (right).

Following the experimental paradigm, the computational framework comprises four stages: H&E image registration, morphology feature extraction, model traing&inference, and label inference. The registration process first aligns all slices to a common coordinate framework (CCF). Subsequently, individual cells are identified and processed through a pathology foundation model for morphology feature extraction. These features are fed into our trained SPONGE for inference, generating predicted spatial multi-omics profiles for slices. Finally, we utilize learned latent embeddings to transfer cell-type and spatial-domain labels from the sequenced slices to the imaged volume. By concatenating the resulting *N* slices, we reconstruct a unified 3D dataset encompassing multi-omics expression, domain architecture, and cell type distribution (Fig. 1a-Output).

The model training of SPONGE and label inference processes are illustrated in Fig. 1b and 1c, respectively. During model training, both the sequenced and their adjacent imaging-only slices (in total 2 *n* slices) were fed into the neural network to optimize the diagonal prediction of omics features and mitigate the staining shift from H&E images (see Fig. S1 and Supplementary Note for staining shift, see Methods for details). For the spatial domain identification, cells were represented using SPONGE latent embeddings, upon which a spatial graph was constructed. A Gaussian Mixture Model was then trained on the anchor slices and applied to the imaging-only slices (see Methods)^7^. In terms of the cell type inference, we take cell type labels from the anchor slices as a reference, and these labels were propagated to the remaining slices based on embedding similarity (see Methods).

In theory, any H&E-based omics prediction methods^6,8-14^ can function as SPONGE by training multiple single-omics prediction models and then inferring on the whole tissue volume. SPONGE distinguishes itself from existing ones by implementing diagonal integration, which inherently facilitates the cross-omics integration during training, thereby enhancing overall model performance. To assess SPONGE in diagonal integration, we implemented extensive assessments (Fig. 1d). Quantitatively, we compared SPONGE with other state-of-the-art methods^6,8-14^ in terms of omics-level Pearson correlation coefficient (PCC, higher better) (Fig. 1d-i,ii,iii) and correlation matrix distance (CMD, lower better) (Fig. 1d-iv) (see Methods). It is apparent that our methods achieve the highest PCC and smallest CMD across all the evaluated omics. In comparison with the second-best methods across the conditions, SPONGE demonstrates performance gains of 66.3%, 46.5%, and 23.5%, respectively. Meanwhile, SPONGE also generated more reliable predictions (the number of high PCC values) over other methods (Fig. S2). Eventually, SPONGE demonstrated an advantage in spatial conservation on the predicted omics layers (Fig. 1d-v)(see Methods).

Qualitative results for slice ID1 regarding predicted Omics 2 and 3 are shown in Fig. 1vi. Compared to the second-best method, mclSTExp^14^, SPONGE generates predictions that more closely resemble the ground truth, particularly in the regions highlighted by the orange and blue boxes (see more in Fig. S3). For cell-type inference, labels were transferred from slices ID10 and ID100 to slice ID1 as a demonstration. The qualitative results are shown in Fig. 1d–vii, with quantitative evaluation presented in Fig. S4a,b. The transferred labels closely match the ground truth, highlighting the effectiveness of the proposed strategy. In Fig. S4c, we further visualize label transfer from anchor slices to their adjacent slices, demonstrating the practical utility of this approach. For spatial domain identification, we compared domain assignments derived from measured spatial multi-omics data with those obtained using our inference strategy (Fig. 1d–viii and Fig. S5a). In addition to visual comparison, four evaluation metrics indicated high consistency of the domain results (Fig. S5b). Overall, these results demonstrate the strong representational capacity of our computational framework and support the reliability of the predicted results for the imaging-only slices.

Following the validation of our multi-omics prediction and label inferences, we reconstructed the 3D architecture of the tissue, incorporating multi-omics profiles, cell-type distributions, and spatial domains. Fig. 2a shows the pathological annotation of the tissue (left) and the 3D distribution of various cell types (right). Domain rendering was performed to create a continuous representation of the scattered cells, which clearly delineates the structure of each region (Fig. 2b). We also examined the gene and protein markers and confirmed their consistency with the regional annotations (Fig. 2c–d, Fig. S6, Fig. S7). Specifically, the proliferation marker Ki67 was highly expressed within the tumor region, while Collagen IV, a basement membrane component, was localized to the boundary. Vascular regions were characterized by high expression of SMA (a vascular smooth muscle cell marker^15^) and CD31(a vascular endothelial marker^16^), and inflammation-associated signatures (*SAA4*^17^, *C6*^18^, and *SOCS3*^19^) were enriched in the peritumoral area, suggesting a localized inflammatory state.

**Fig. 2.**
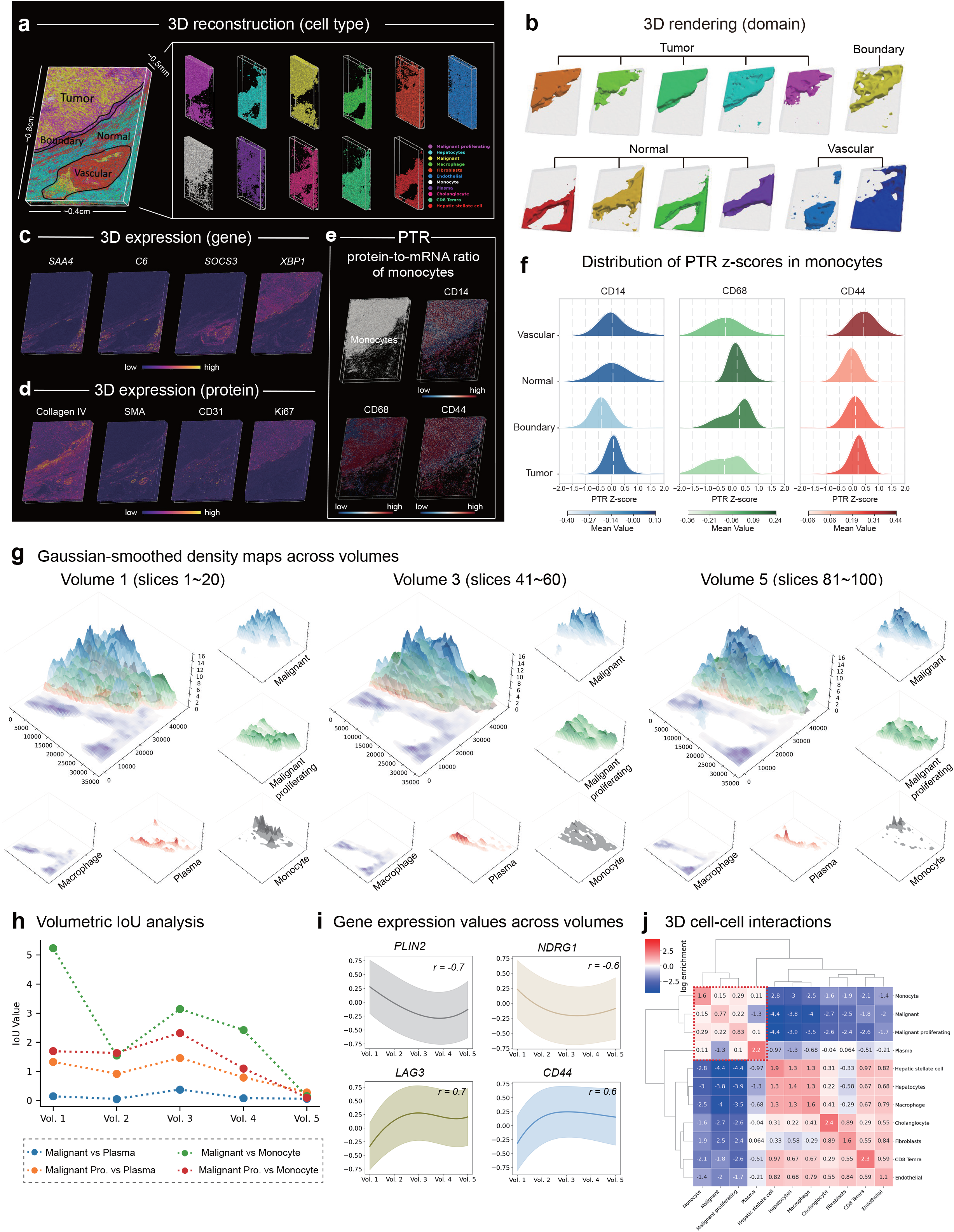
Downstream analysis on the generated 3D spatial multi-omics data. a, The left panel displays an integrated view of all data points with pathologist-defined regions and physical tissue dimensions. The right panel isolates distinct cell types by color. b, 3D renderings of individual spatial domains, grouped by pathological annotations. c-d, 3D visualizations showing the spatial distribution of selected gene (c) and protein (d) signatures. e, Spatial distribution of monocytes and three protein-to-RNA ratio (PTR) scores. PTR acts as a proxy for translational efficiency variance, where positive values indicate a ratio above the population average. f, Quantification of the monocyte PTR scores (CD14, CD68, CD44) mapped across four pathological regions (tumor, boundary, normal, and vascular). Kernel density curves highlight the median (white lines) and mean (color). g, Illustrations of the gaussian-smoothed density maps of individual cell types and their overlap across volumes. Intended cell types include malignant, malignant proliferating, monocyte, plasma, and macrophage. h, Volumetric IoU values of malignant and malignant proliferation with plasma and monocyte. i, Expression values of monocyte-status-related marker genes across volumes. j, The heatmap of the log enrichment regarding 3D cell–cell interactions across different cell types, with hierarchical clustering revealing distinct interaction patterns. Monocytes and malignant cell populations exhibit notably reduced interactions with most other cell types, while stromal and epithelial cells display stronger mutual connectivity. The tumor region was indicated with a dashed red box.

Having established the accurate diagonal integration of multi-omics features, we further explored the protein-to-mRNA ratio (PTR) as a proxy for translational efficiency in 3D data volume (see Methods). Given the observed dense spatial colocalization between monocytes and malignant cells (Fig. 2a), we particularly investigated the spatial dynamics of PTR within the monocyte population (Fig. 2e). Quantitative and qualitative analysis revealed that translation rates for key monocyte proteins vary across tissue domains. For example, the canonical monocyte marker CD14 exhibited elevated PTR scores in normal regions compared to other areas. For example, the PTR score of the canonical monocyte marker CD14 is reduced in the boundary region and relatively higher in other regions. CD44 shows higher PTR scores in vascular and tumor regions, whereas CD68 scores are lower in these regions (Fig. 2f). These spatial disparities in PTR scores suggest region-specific alterations in monocyte states within the tumor microenvironment, potentially reflecting its functional heterogeneity across regions.

To capture biological transitions along the z-axis, we generated Gaussian-smoothed density maps for key cellular populations (malignant, malignant proliferating, monocyte, plasma, and macrophage) across volumes (every 20 slices as a volume) in Fig. 2g and Fig. S8. We observed spatial segregation between malignant cells and macrophages by intervening plasma cells, contrasted by dense colocalization between monocytes and malignant populations. Volumetric Intersection over Union (IoU) analysis quantified a progressive attenuation of monocyte-malignant co-localization along the z-axis (Fig. 2h). This spatial decoupling suggests a niche-specific transition from inflammatory signaling toward mature macrophage phenotypes in deeper tumor regions. This was further characterized by the expression of four marker genes: the down-regulation of metabolic and hypoxic stress markers (*PLIN2, NDRG1*) and the concomitant up-regulation of immune checkpoint and adhesion regulators (*LAG3, CD44*) in Fig. 2i. Together, these shifts mark a terminal transition from acute stress adaptation toward an immunosuppressive, tumor-associated macrophage (TAM) phenotype^20^.

Finally, we employed permutation tests to identify enriched 3D cell-cell interactions across the tissue volume (Fig. 2j). Clustering analysis partitioned the tissue into distinct tumor and non-tumor domains. Within the latter, 3D spatial mapping revealed significant co-localization of hepatic stellate cells and macrophages, likely facilitating local extracellular matrix remodeling and supporting residual hepatocytes. The pronounced spatial segregation between this stromal cluster and the malignant core suggests a structural barrier that may impede immune infiltration into the tumor nests.

In summary, we have presented Histo3D-MO, a hybrid pipeline for single-cell-resolution 3D spatial multi-omics synthesis. This holds implications for practical studies, addressing a long-standing technical void due to the inherent challenges. Histo3D-MO overcomes these limitations by leveraging diagonal single-omics data alongside dense H&E histology images to enable economical 3D multi-omics mapping. A key step of Histo3D-MO is the use of the SPONGE framework, which leverages H&E-derived morphological features to facilitate flexible multi-omics diagonal integration. Through the implementation of specialized cell-type and spatial-domain inference strategies, we have demonstrated their widespread utilities across scenes. These strategies not only offer granular insights into the transcript–protein relationship and spatial architecture but also provide a scalable approach to reconstructing complex molecular environments for complex biological analysis.

While Histo3D-MO provides a robust framework for 3D spatial multi-omics reconstruction, certain limitations warrant consideration. First, the reliance on serial sectioning is inherently labor-intensive and introduces the potential for tissue loss or damage during sample preparation. Furthermore, the image registration process may introduce subtle alignment artifacts, which could subsequently impact the overall fidelity of the 3D volumetric reconstruction. Ultimately, despite these limitations, Histo3D-MO represents a pioneering step toward resolving this grand challenge, establishing a powerful new paradigm for spatial biology.

## 2. Methods

### Experimental Paradigm

The Histo3D-MO pipeline begins with rigorous data preparation. The intact tissue block undergoes serial sectioning along the z-axis, with slice thicknesses optimized for specific experiments, e.g., 5∼10 µm for Xenium spatial sequencing and approximately 5 µm for H&E imaging. While the entire cohort of *N* slices is subjected to H&E imaging, a representative subset of *n* sparse slices is selected for distinct spatial single-omics sequencing. To maximize cost-efficient 3D spatial multi-omics mapping, the *n* slices are profiled without omics overlapping. The selection of this subset is strategically designed to maximize volumetric representation, facilitating the downstream prediction of molecular profiles from morphological data. Hereby, the boundary (first and last) slices are prioritized for sequencing given the large morphology and omics variations. Ultimately, this paradigm produces a comprehensive dataset containing a continuous morphological H&E sequence with discrete *n* spatial single-omics measurements.

### The in-house Spatial Multi-omics Hepatocellular Carcinoma (moHCC) Dataset

We applied our experimental paradigm on an intact hepatocellular carcinoma (HCC) tissue block (Ethics approval number: 2024-R318). The tissue block was serially sectioned into a total of 100 slices (*N* =100). Among these, slices ID1, ID10, and ID100 (*n*=3) were selected for single-cell-resolution spatial multi-omics profiling. Specifically, each selected slice was subjected to Xenium Prime 5K for large-scale spatial transcriptomic and PhenoCycler-Fusion (PCF) for spatial proteomic profiling (see Supplementary Note for details). To simulate a multi-modal environment for downstream diagonal integration evaluation, we treated each slice as a distinct omics layer paired with its respective morphological image. Specifically, we established non-overlapping transcriptomic panels by pooling the top 500 highly variable genes (HVGs) from each slice into a combined set of 569 genes. This consolidated set was then bifurcated, assigning 284 genes (Panel A) to slice ID1 and 285 genes (Panel B) to slice ID10. Finally, slice ID100 was designated for spatial proteomics using a 21-protein panel, successfully generating a heterogeneous dataset for robust model validation.

### Cell Segmentation for Imaging-only Slices

To identify single-cell structures from H&E images, we applied the deep learning–based segmentation framework Cellpose^22^ (v3, pretrained cyto3 model) for automated segmentation. Cellpose is built upon a U-Net architecture and performs instance segmentation by predicting pixel-wise spatial gradient flows, enabling robust performance across diverse tissue types and imaging conditions. To facilitate efficient processing of large whole-slide images, we followed the “Big Data” workflow provided in the Cellpose tutorial (the cellpose.contrib.distributed_segmentation module). GPU acceleration was enabled for inference. The model type was set to cyto3, with the target cell diameter specified as 30. A single-channel input configuration was used (channels = [0, 0]), and 3D segmentation was disabled (do_3D = False). The block size for distributed processing was set to 4096 × 4096 pixels.

Segmentation outputs consisted of instance label masks, in which each pixel was assigned a unique cell. The resulting masks were stored as NumPy arrays and archived in the seg.npy format to ensure reproducibility and facilitate downstream analyses. To obtain spatial coordinates for individual cells, we computed the centroid of each instance based on the segmentation masks. Specifically, centroids were calculated using the scipy.ndimage.center_of_mass function from SciPy. The resulting two-dimensional centroid coordinates were exported as CSV files for subsequent spatial analyses.

### Cell Modeling and Hierarchical Feature Extraction

Subsequent to registration and segmentation, individual cells within each tissue slice were modeled to capture detailed biological morphology. While we utilized the provided cell masks for sequenced slices, our custom segmentation pipeline was applied to the imaging-only slices (see above). Using the resulting cell-centric coordinates, we extracted 128 × 128 pixel patches (approximately 30µm) to represent each cell. Given a mean cell diameter of ∼10µm, this sampling strategy ensures that each patch encapsulates both the target cell and its immediate spatial neighbors, thereby preserving critical microenvironmental context.

To capture multi-scale biological signals, we utilized the pre-trained UNI^23^ model within a hierarchical extraction strategy. Patches were reshaped to 256 × 256 and then computed by UNI as *f*_map_ ∈ ℝ^1024×26×26^. We then extracted two distinct scales of information: a local representation *f*_local_, computed by average pooling the central 4 × 4 grid corresponding to the target cell; a global representation *f*_global_, derived from the average pooling of global features. The final representation *f* ∈ ℝ ^2048^ consists of the concatenated local and global vectors, ensuring the model accounts for both individual cell characteristics and the surrounding histopathological environment (see Fig. S9a).

#### Domain identification for sequenced slices

For the sequenced slices, we first generate latent embeddings for all cells, denoted as 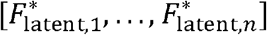, which are concatenated to form the global embedding matrix 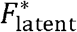. These embeddings undergo z-score standardization and Principal Component Analysis (PCA) to produce a reduced-dimension feature set, 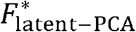 . To incorporate spatial context, both 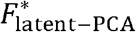 and the corresponding cellular spatial coordinates are integrated into the CellCharter framework^7^. This process encompasses spatial graph construction, neighborhood feature aggregation, and the training of a Gaussian Mixture Model (GMM) to delineate distinct tissue domains.

#### Generalization to Imaging-Only Slices

To extend these domains to imaging-only slices, we generate a latent embedding 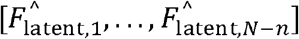, concatenated as 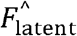 . To maintain cross-slice consistency, 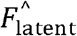 is pre-processed using the standardization parameters and PCA transformation matrix derived from the initial sequenced (training) slices, yielding 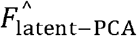 . These refined features, alongside their spatial coordinates, are subsequently processed by the pre-trained CellCharter model for automated domain assignment. This hierarchical protocol ensures high identification accuracy and cross-modal alignment while significantly reducing the computational overhead typically associated with iterative clustering methods.

### Cell Type Inference

Characterizing cellular identities is fundamental to deciphering tissue architecture; however, transferring established cell-type labels from sequenced reference data to imaging-only slices presents a substantial computational challenge. To address this, we developed a robust retrieval-based strategy that leverages the high-fidelity latent embeddings generated by the SPONGE model.

Specifically, the process utilizes the sequenced latent embeddings 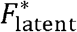 and their corresponding validated cell-type labels Ct* as a reference library. For each individual cell in the imaging-only slices, we compute the pairwise embedding distance between its specific latent representation 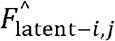 (where *i* and *j* denote the slice and cell IDs, respectively) and the reference matrix 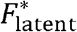 . To ensure the reliability of the transfer, we implement a majority voting strategy based on the top-*k* nearest neighbors in the reference library. A cell-type label is assigned only if a single type constitutes a strict consensus (exceeding *k*/2) among the top- *k* neighbors. Cells failing to meet this confidence threshold are excluded from the analysis to mitigate classification uncertainty. This stringent filtering protocol ensures that the resulting spatial maps represent high-confidence biological identities.

### 3D Cell-cell Interaction Analysis

To characterize the spatial architecture and potential intercellular interactions in the 3D view, we represented the tissue organization as a proximity graph derived through Delaunay triangulation. This topological framework enabled the construction of a spatial adjacency matrix to quantify interactions across diverse cell populations. To distinguish biologically meaningful associations from stochastic noise, we performed a permutation test by iteratively shuffling cell-type labels to establish a null distribution. This approach allowed us to identify statistically significant nearest-neighbor preferences and quantify the enrichment or depletion of specific cell-type pairings. These organized spatial relationships were visualized via heatmaps, revealing the recurring structural motifs that define the cellular microenvironment.

### Pathological Annotation

Pathological region labels were manually annotated by two expert pathologists based on H&E images. Based on the histological features, the tissue was categorized into four primary regions: tumor, boundary, normal, and vascular. These expert-annotated labels were subsequently used for 3D visualization and downstream biological analyses.

### 3D Rendering

After completing the 3D reconstruction, we utilized the “3D rendering” module implemented in Spateo^28^ to transform discrete spatial points into a continuous representation for visualization. This module is primarily built upon the PyVista framework. Following the official Spateo tutorial, we implemented the workflow using functions such as construct_pc and construct_surface to generate the corresponding representations. The spatial omics data in .h5ad format were subsequently converted into .vtk files for model storage, and the resulting models were visualized using the Plotter class from the PyVista package. We also adjusted various parameters according to the Spateo guidelines, such as the camera position settings (the cpo parameter), to optimize visualization quality. Specifically, the domain label was selected as the rendering subset, and each cluster was rendered individually. The final results were then organized and presented according to the pathological annotation categories.

### PTR analysis

To characterize the relative abundance relationship between protein and transcript at single-cell resolution while minimizing distributional differences between molecular layers, we defined a Protein-to-Transcript Ratio (PTR) score to quantify each matched protein–RNA pair in every cell. First, proteins were matched to their corresponding RNA features, and only successfully paired protein–RNA combinations were retained for downstream analysis.

For each cell *j* and each matched pair *i*, protein and RNA expression matrices were independently normalized by library size scaling to a fixed total of 10,000 counts per cell:

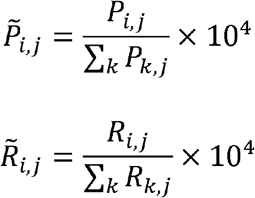

where 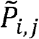 and 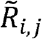 denote the protein and RNA counts, respectively. We then defined the raw protein-to-transcript ratio (rPTR) as the log2-transformed ratio:

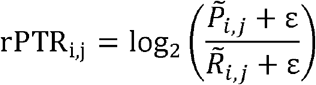

where ε = 10^−10^ was introduced to avoid numerical instability due to zero values. Because different genes exhibit substantial variation in overall abundance and dynamic range, direct comparison of raw PTR values may be confounded by gene-specific distributions. Therefore, for each matched pair, raw PTR values were standardized across all cells:

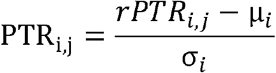

where *µ*_*i*_ and σ_*i*_ represent the mean and standard deviation of rPTR across all cells for matched pair *i*.

The standardized PTR thus reflects the deviation of a given cell from the global distribution of that protein–RNA pair. Positive values indicate that the protein-to-transcript ratio is higher than the population average, whereas negative values indicate a lower-than-average ratio.

Importantly, because protein and RNA measurements originate from distinct molecular layers and are subject to different detection efficiencies, degradation rates, and technical noise, PTR score should not be interpreted as a strict translation rate or an exact molecular stoichiometric ratio. Instead, it provides an approximate measure of relative protein-to-transcript variation across cells, capturing potential differences in post-transcriptional regulation. Consequently, PTR is more appropriate for comparative analyses across cells or spatial regions than for absolute quantification of translational kinetics.

### Evaluation Metrics

To rigorously quantify the performance of our omics prediction framework, we employed three complementary evaluation metrics that capture different aspects of statistical and structural similarity between predicted and ground-truth expression profiles.

#### Pearson Correlation Coefficient (PCC)

The PCC was utilized to measure the linear relationship between predicted and observed molecular intensities. It provides a direct assessment of how well the model reconstructs the absolute magnitude of gene or protein expression levels across spatial locations. PCC is defined as:

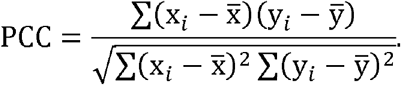

#### Spearman Correlation Coefficient (SPCC)

The SPCC assesses the monotonic relationship between predicted and observed expression levels by calculating the Pearson correlation of their respective ranks. For a set of *n* observations, let rg(*x*_*i*_) be the rank of the *i*-th observed value and rg(*y*_*i*_) be the rank of the *i*-th predicted value. The coefficient is formulated as:

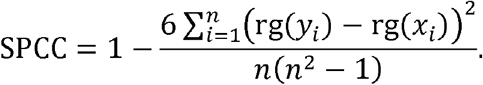

This metric is particularly valuable for omics data, as it is less sensitive to outliers and does not assume a strictly linear relationship between the modalities.

#### Correlation Matrix Distance (CMD)

The CMD is a measure of the distance between two correlation matrices, providing a global assessment of how well the model preserves the co-expression structure (the “inter-omic” dependencies). Let R be the ground-truth correlation matrix and 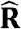 be the predicted correlation matrix. The CMD is defined as:

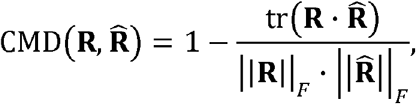

where tr(·) denotes the trace of the product of the matrices and ||·||_*F*_ denotes the Frobenius norm of the matrix. The CMD ranges from 0 to 1, where a value of 0 indicates that the two correlation matrices are identical (perfect preservation of the regulatory network), and a value of 1 indicates they are maximally dissimilar.

#### Spatial conservation analysis

To evaluate whether the predicted molecular features preserve the spatial organization of the measured data, we performed a spatial conservation analysis based on spatial autocorrelation and feature similarity. Specifically, we first computed the spatial autocorrelation statistic Moran’s I for each measured and predicted omics feature using the spatial coordinates of individual cells. Moran’s I quantifies the degree of spatial clustering of molecular signals, enabling assessment of whether the predicted features retain the intrinsic spatial patterns present in the measured data.

Next, to quantify the agreement between measured and predicted spatial patterns, we calculated the PCC between the Moran’s I values derived from measured and predicted features. The correlation analysis was performed separately for each omics panel to ensure panel-specific evaluation of spatial structure preservation. A higher PCC indicates stronger consistency in spatial autocorrelation between measured and predicted features, suggesting that the model effectively preserves spatial molecular organization.

## Acknowledgement

This study was supported by the Computational Biology Program (no. 25JS2850200, to Z.Y.) of Science and Technology Commission of Shanghai Municipality (STCSM), National Natural Science Foundation of China (nos. 62303119 and 32470706 to Z.Y.), Shanghai Science and Technology Development Funds (no. 23YF1403000, to Z.Y.), Fund of Fudan University and Cao’ejiang Basic Research (no. 24FCA10, to Z.Y.).

## Author Contributions Statement

Z.Y. conceived and designed the study. Z.W., Y.Y., X.Y. developed the computational methods, performed the analysis, and wrote the manuscript. D.Z., and Q.Z. collected and processed the datasets. Y.D. and Z.H. polished the paper writing and designed the figures.

## Data Availability

The dataset will be released after acceptance.

## Code Availability

The Pytorch implementation of this study is available at https://github.com/ZacharyWang-007/Histo3D-MO.

